# Structural defects in amyloid-β fibrils drive secondary nucleation

**DOI:** 10.1101/2025.03.31.646415

**Authors:** Jing Hu, Tom Scheidt, Dev Thacker, Emil Axell, Elin Stemme, Urszula Łapińska, Stefan Wennmalm, Georg Meisl, Samo Curk, Maria Andreasen, Michele Vendruscolo, Paolo Arosio, Andela Saric, Jeremy D. Schmit, Tuomas P. J. Knowles, Emma Sparr, Sara Linse, Thomas C. T. Michaels, Alexander J. Dear

## Abstract

The nucleation of amyloid fibrils from monomeric protein, catalyzed by the surface of existing fibrils, is an important driver of many disorders such as Alzheimer’s and Parkinson’s diseases. The structural basis of this secondary nucleation process, however, is poorly understood. Here, we ask whether secondary nucleation sites are found predominantly at rare *growth defects*: defects in the fibril core structure generated during their original assembly. We first demonstrate using the specific inhibitor of secondary nucleation, Brichos, that secondary nucleation sites on Alzheimer’s disease-associated fibrils composed of Aβ40 and Aβ42 peptides are rare compared to the number of protein molecules they contain. We then grow Aβ40 fibrils under conditions designed to eliminate most growth defects while leaving the regular fibril morphology unchanged, and confirm the latter using cryo-electron microscopy. We measure both the ability of these *annealed* fibrils to promote secondary nucleation and the stoichiometry of their secondary nucleation sites, finding that both are greatly reduced as predicted. Re-analysis of published data for other proteins suggests that fibril growth defects that expose monomer planes or other structural units may also drive secondary nucleation generally, across most or all amyloids. These findings could unlock structure-based drug design of therapeutics that aim to halt amyloid disorders by inhibiting secondary nucleation sites.

## 1 Introduction

Amyloid fibrils are characterized by the stacking of proteins into β-sheet-rich aggregates with a hydrophobic core. They can serve functional purposes but are also frequently associated with pathology [1]. Formation of pathological amyloids from amyloid-β(1-42) peptide (Aβ42), amyloid-β(1-40) peptide (Aβ40) and tau protein are key events in Alzheimer’s disease (AD) [2–4]. The spontaneous formation of amyloid fibrils requires a minimum of two reaction steps: a slow primary nucleation process producing very short new fibrils from monomeric protein, and the rapid elongation of fibrils by addition of monomer to their ends. [5] Often, an additional process called secondary nucleation is also present, in which new fibrils are formed by monomer association on the surface of existing fibrils [6–9]. Amyloidogenic proteins whose aggregation has been found to depend on secondary nucleation includes type II diabetes-associated islet amyloid polypeptide (IAPP) [10, 11], Parkinson’s disease-associated *α*-synuclein [12], AD-associated tau [13], and length variants of the A*β* peptide [14, 15]. Once a critical concentration of aggregates has formed in these systems, the vast majority of new fibrils do not form through primary nucleation but rather via secondary nucleation [15, 16]. Rapid secondary nucleation is characteristic of many disease-associated amyloids, due both to its auto-catalytic effect on fibril proliferation and to its ability to produce large quantities of toxic protein oligomers [9, 17].

The fundamental importance of secondary nucleation extends beyond amyloid chemistry. It is known to be present for example in the solution-to-solid phase transition of a number of crystalline materials [18–20], and in the aggregation of the mutant haemoglobin HbS linked to sickle-cell anaemia [8, 16, 21, 22]. However, despite this importance, exactly how fibril surfaces catalyze nucleation is still not known. In particular, it is not clear where on fibrils secondary nucleation occurs, and the nature of the catalytic site. Numerous possibilities have been hypothesized; a common assumption is that catalysis is uniform across the entire fibril surface, or at least that catalytic sites occur at every plane in the fibril (see Fig. 1**b****(i)**). Nonetheless, several recent studies cast doubt on this hypothesis. It was inferred in ref. [23] that secondary nucleation of Aβ42 occurs at isolated sites on the fibril that are separated by inactive surface. It was furthermore indirectly estimated that these sites must be relatively rare, with stoichiometry no higher than 1 site per 30 protein monomers in the fibril, and several other studies support the rarity of Aβ secondary nucleation sites [24–27]. However, the stoichiometry of these sites has never been quantified directly.

**Fig. 1.**
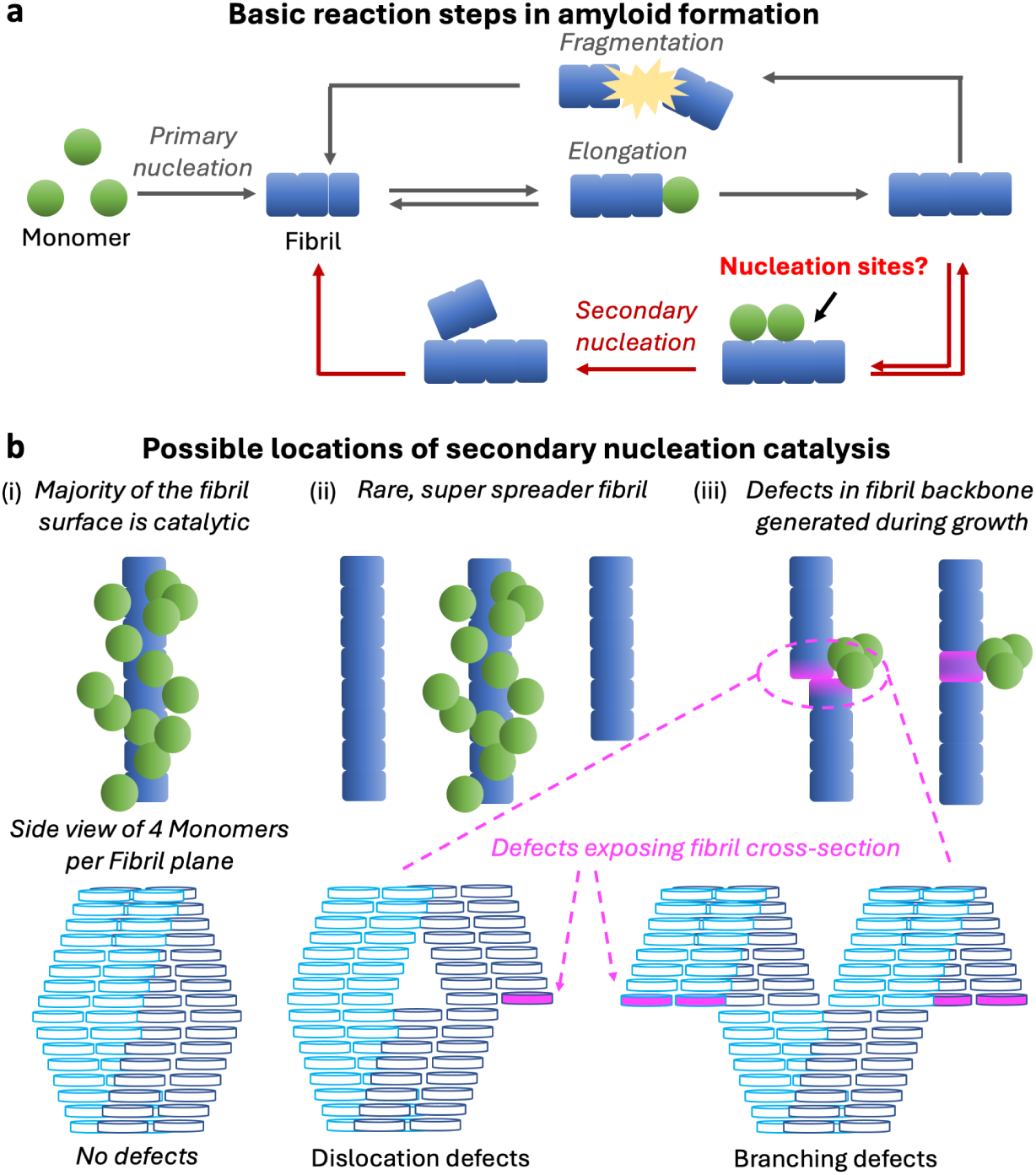
Secondary nucleation is a core process driving amyloid fibril proliferation; however, its structural basis is not understood. **a**: Overall reaction network governing amyloid fibril formation. Secondary nucleation is often responsible for most new fibril formation. **b**: Possible ways amyloid fibrils might promote secondary nucleation of new fibrils (and oligomers). **(i)**: Uniform catalysis of nucleation across the entire fibril surface. **(ii)**: Rare “super-spreader” fibrils have dense catalytic sites, with most fibrils being incapable of promoting secondary nucleation. **(iii)**: Secondary nucleation sites are defects in the fibril core, created during fibril elongation. Such defects can coincide with dislocations or branches (LHS, and bottom row), although non-offset defect structures are also possible (RHS). Fibril cross-sections denoted by purple units and marked by arrows are exposed at dislocation or branching defect sites. The fraction of monomer units exposed may differ from that illustrated here. For simplicity, defects in subsequent figures will be represented schematically as dislocations only.

Another proposed explanation for substoichiometric secondary nucleation sites is that catalysis is localized at rare “super-spreader” fibrils with dense secondary nucleation sites of the same periodicity as the fibrils, with the remaining majority of fibrils being incapable of secondary nucleation [28] (see Fig. 1**b****(ii)**). However, this would be inconsistent with isolated Aβ42 secondary nucleation sites. Moreover, recent studies have found that most Aβ42 fibrils have comparable catalytic activity [28], and that almost all Aβ42 fibrils possess non-periodic secondary nucleation sites [29]. In light of the above findings, it has been proposed that these sites instead correspond to structural defects. The lack of relationship detected between occurrence of deformed or curved Aβ42 fibrils and secondary nucleation rate would appear to rule out that these defects are a consequence of mechanical strain [28].

Instead, defects in the fibril core are a more plausible candidate [23]. Those types of defect structure that permit the next fibril plane to be assembled with the correct structure on top of the defective fibril plane without imposing an insurmountable free energy penalty should exist throughout the fibril length coexisting with otherwise uniform fibril morphology. One example of such a defect is when the monomers in a new fibril plane bind correctly with respect to one another and with the correct fold but laterally offset and potentially also out-of-register from the monomers in the preceding fibril plane during fibril elongation [30, 31]. Because the next fibril plane can be assembled in perfect alignment with the offset plane, the free energy penalty for doing so is minimal (see Fig. 1**b****(iii)**). Small dislocations only partially exposing a filament’s cross-section are possible, as are larger dislocations that lead to entire filaments being misaligned, or even to fibril branching. Other plausible defect structures include (but are not limited to) fibril planes in which one or more of the monomers is only partially correctly folded, exposing part of the fibril’s hydrophobic core; however, such defects may not be stable enough to appear with significant frequency in fibrils.

Crucially, the hypothesis that growth defects cause secondary nucleation cannot be directly tested using structural data such as cryo-electron microscopy or solid-state nuclear magnetic resonance, which report only periodic or averaged properties of fibrils. Here, we set out to determine the identity and origin of amyloid-β secondary nucleation sites *in vitro*. We first quantify the stoichiometry of Aβ42 and Aβ40 secondary nucleation sites using two distinct experimental methods, finding them to occur roughly only every 150 or every 50 monomers in the fibrils, respectively. We then grow Aβ40 fibrils under conditions designed to eliminate most growth defects, and show using fluorescence correlation spectroscopy that their secondary nucleation site stoichiometry is also greatly reduced. We use kinetic assays to demonstrate that the secondary nucleation rate of such *annealed* fibrils is reduced commensurately. High resolution cryo-electron microscopy finds no evidence of difference in morphology between normal and annealed fibrils, consistent with the large difference in their secondary nucleation site frequency and rates being caused by differences in defects. We estimate the size of the Aβ defects by examining their stoichiometry in the absence of supersaturation, finding them to incur a free energy penalty of around 50% of the binding energy of a single monomer in the correct conformation. Based on thermodynamic arguments and re-analysis of published studies, we finally hypothesize that defect-based secondary nucleation is not solely a feature of Aβ but a general phenomenon in amyloid formation.

## 2 Results

### 2.1 Brichos binds A*β*42 amyloid fibrils tightly with low stoichiometry

We first sought to characterise the interaction between the Brichos chaperone domain and Aβ42 fibrils and quantify the absolute amount of chaperone bound to fibrils. To this end, we employed a microfluidic diffusional sizing (MDS) platform that was previously developed to quantify the interactions between biomolecules in solution under native conditions [32, 33]. In brief, this technique utilises the diffusion profiles of molecules in solution collected at multiple positions as they flow through a microfluidics channel (Fig. 2**a**). From the analysis of these diffusion profiles using the advection-diffusion equation it is possible to extract the distribution of diffusion coefficients. Brichos molecules that are bound to amyloid fibrils diffuse through the channel cross-section much more slowly than unbound Brichos molecules, exhibiting distinct diffusion profiles that can be detected [33]. Using this platform we quantified Alexa- 488 labelled proSP-C Brichos binding to Aβ42 amyloid fibrils by incubating 24 µM of fibrils in the presence of Brichos with increasing concentrations at 21*^◦^*C.

**Fig. 2.**
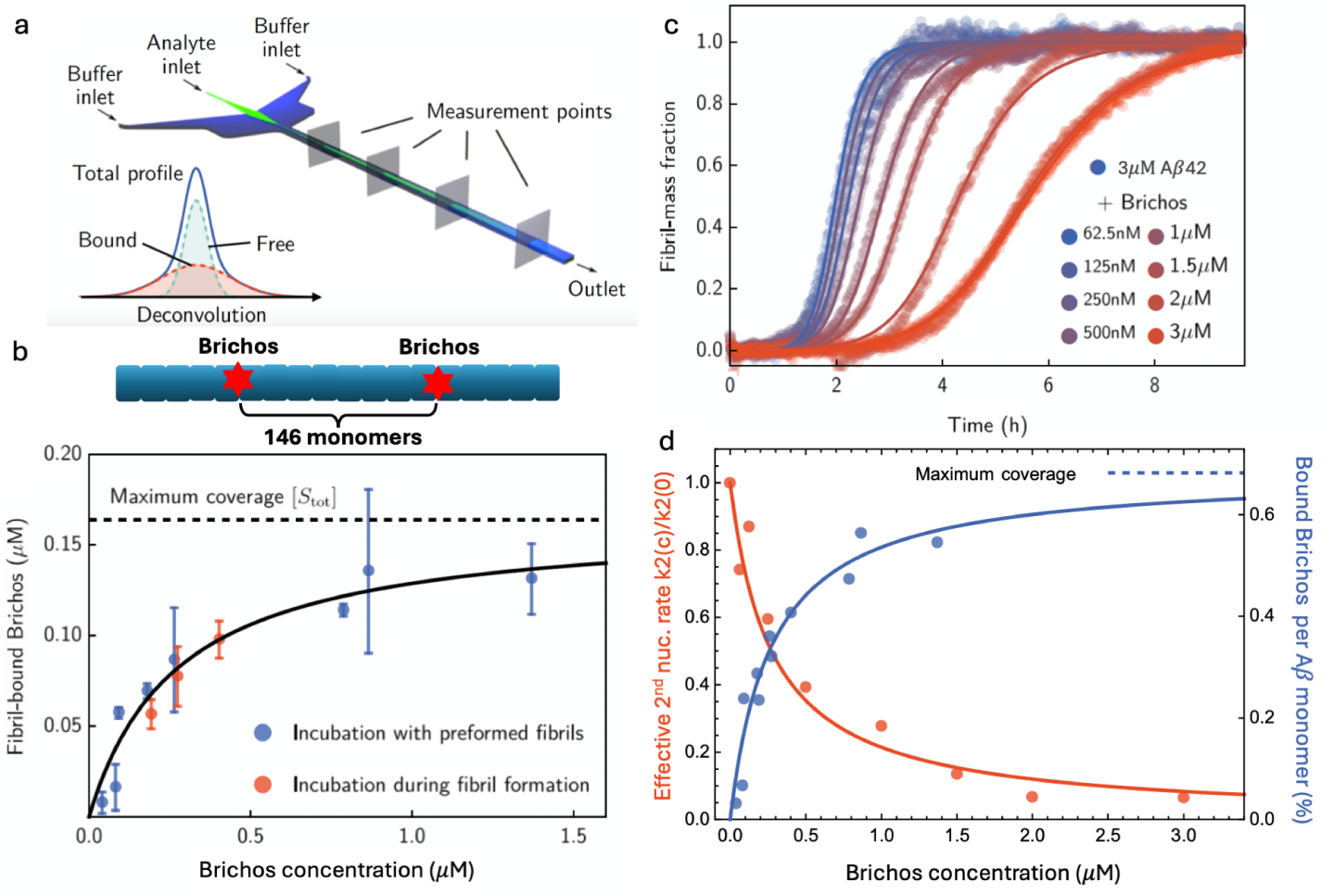
Stoichiometry of Brichos binding to Aβ42 fibrils is low, and is predictive of secondary nucleation inhibition. **a**: Illustration of MDS experiment: free chaperone molecules diffuse rapidly to the edges of the channel, while chaperone molecules that interact with amyloid fibrils display slower lateral diffusion and remain therefore localised in the centre of the channel cross-section. Fibril-bound chaperone concentration can be inferred by mathematical analysis of the diffusion profile. **b**: Binding of Alexa-488 labelled proSP-C Brichos to 24 µM Aβ42 fibrils. Error bars are standard deviations calculated from 2 (blue) or 3 (red) repeats. The regression line represents the best fit to the Langmuir adsorption isotherm with *K_D_* = 273 nM. Binding stoichiometry: maximum coverage corresponds to one Brichos molecule bound per ca. 146 Aβ42 monomers. **c**: Kinetic aggregation profiles of 3 µM Aβ42 with 1% seed in the presence of increasing concentrations *C* of proSP-C Brichos. The solid lines indicate global fits using a rate law for Aβ42 in the presence of a secondary nucleation inhibitor that includes an effective secondary nucleation rate constant *k*_2_(*C*) (see SI Sec. 2). **d**: Decrease in effective secondary nucleation rate constant, *k*_2_(*C*)*/k*_2_(0), with increasing Brichos concentration *C* (red data, left axis), overlaid with the chaperone binding curve to fibrils (blue, right axis). The solid red line indicates the predicted decrease in secondary nucleation rate constant *k*_2_(*C*)*/k*_2_(0) = 1*/*(1 + *C/K_D_*) if the secondary nucleation sites are the same as the Brichos binding sites; this model fits the data very well. All experiments in this figure are done in 20 mM sodium phosphate, 0.2 mM EDTA at pH 8.0 and 21*^◦^*C.

Two different incubation procedures were followed: in a first approach, we measured the binding of Brichos to Aβ42 fibrils generated in the absence of the molecular chaperone. In a second experiment, Brichos was incubated together with monomeric Aβ42 at 37*^◦^*C until essentially all monomers had converted into fibrils, followed by 48 h incubation at 21℃, after which the binding was evaluated. The resulting binding curves are shown in Fig. 2**b**; the data obtained through the first and second procedures (red and blue points) are closely similar.

To obtain quantitative information about the binding process we fit to the experimental data the Langmuir adsorption isotherm [34] 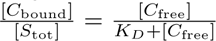 is the dissociation constant, [*C*_bound_] and [*C*_free_] are the concentrations of the bound and free chaperone, respectively, evaluated by the size distributions measured by the microfluidics diffusion technique. [*S*_tot_] is the total concentration of binding sites available on a Aβ42 fibril surface, representing the maximum concentration of bound molecular chaperone. This procedure yields a dissociation constant of *K_D_* = 273 nM at 21*^◦^*C and a maximal concentration of bound chaperone of [*S*_tot_] = 164 nM. Dividing [*S*_tot_] by the total concentration of monomer equivalents of Aβ42 in the fibrils reveals that maximum coverage corresponds to a binding stoichiometry *s* of about one Brichos molecule per (24 µM)/(164 nM)=146 Aβ42 monomers (Fig. 2**b**), i.e. *s* = 1*/*146.

### 2.2 Modulation of A*β*42 aggregation by Brichos reveals rare secondary nucleation sites

We next sought to relate our binding measurements to the ability of Brichos to inhibit secondary nucleation of Aβ42 fibrils. To this end, we incubated 3 µM of Aβ42 at 21*^◦^*C and monitored the time course of amyloid fibril formation by ThT fluorescence in the absence and presence of increasing concentrations of Brichos (Fig. 2**c**). We observe that the addition of Brichos delays aggregation by an amount that depends on the concentration of the chaperone, in line with previous reports [24]. To disentangle the effect of Brichos on the rate constants for the individual microscopic steps of aggregation (primary nucleation *k_n_*, secondary nucleation *k*_2_ and elongation *k*_+_), we compared the experimental kinetic profiles with a theoretical chemical kinetics model of aggregation in the presence of Brichos (see SI Sec. 2). In this model chaperone molecules can bind surface sites of fibrils and in this manner reduce the rate of secondary nucleation, while leaving the rate of the other aggregation steps unaffected [35]. A prediction from this model based on the unperturbed aggregation rate parameters obtained in the absence of Brichos (leftmost curve in Fig. 2**c**) and the rate parameters determined from the binding experiment shows good agreement with the experimental data. A comparison between the effective rate constant for secondary nucleation *k*_2_ with increasing Brichos concentration and the binding curve of Brichos to fibrils confirms that the extent of inhibition of Aβ42 by Brichos correlates with the amount of chaperone bound to fibrils (Fig. 2**d**). Strikingly, we see that almost complete suppression of secondary nucleation (reduction of *k*_2_ by *>* 90%, see 2 and 3 µM Brichos datapoints) corresponds to *>* 90% of the maximal Brichos coverage [*S*_tot_]. We therefore obtain sub-stoichiometric inhibition of secondary nucleation by Brichos, with almost complete suppression of secondary nucleation obtained with only one fibril-bound chaperone molecule per about 150 monomers. These results imply that Brichos binding sites and secondary nucleation sites are one and the same. Under these fibril formation conditions, secondary nucleation sites therefore have a stoichiometry of *s* = 1*/*146, i.e. a frequency of approximately 1 site per 150 monomers in the fibril. With four monomers per plane with two monomers per filament plane and two such filaments in an A*β*_42_ fibril [36, 37], this stoichiometry corresponds to one catalytic site per 36 fibril planes on average. A recent evaluation of the density of fibril protrusion during an ongoing Aβ42 aggregation process at pH 6.8 found a distribution peaking at around 1 protrusion per 28 planes, or per 112 monomers, i.e. similar to the rarity of secondary nucleation sites found here at pH 8.0 [38].

### 2.3 Fibrils possessing fewer growth defects can be generated without measurably altering morphology

The low stoichiometry of Aβ42 secondary nucleation sites we have found in this study supports the hypothesis that these sites are rare growth defects in the fibril core structure. To verify this hypothesis, we sought to produce Aβ fibrils under conditions designed to largely eliminate growth defects, so we could subsequently test whether their secondary nucleation sites are also eliminated.

Near thermodynamic equilibrium, the stoichiometry of growth defects (the number of defects per monomer in the fibril) should follow the Boltzmann distribution, becoming the exponential of the free energy penalty of forming a defect. Because fibrils are highly stable, growth defects are expected to incur large free energy penalties. The exponential dependence therefore strongly suppresses the occupancy of growth defect states, leading to low rates of incorporation of growth defects into the fibril. However, kinetic experiments typically rely on high initial supersaturation (i.e. monomer concentration) to ensure adequate fibril yield and sufficiently fast aggregation reactions. Fibril elongation and its reverse, dissociation, are consequently far from equilibrium during the aggregation process, causing rapid net growth of fibrils. The average time between successive monomer binding events becomes much smaller than the average time taken for an incorrectly bound monomer to detach [31]. Therefore, many fibril planes are bound sequentially on top of a defective fibril plane before it can disassemble. The resultant very strong kinetic trapping of defective fibril planes and reduction in defect dissociation in the growing fibril (Fig. 3**a**) is expected to increase defect stoichiometry [31, 40] (see also SI Sec. 6).

**Fig. 3.**
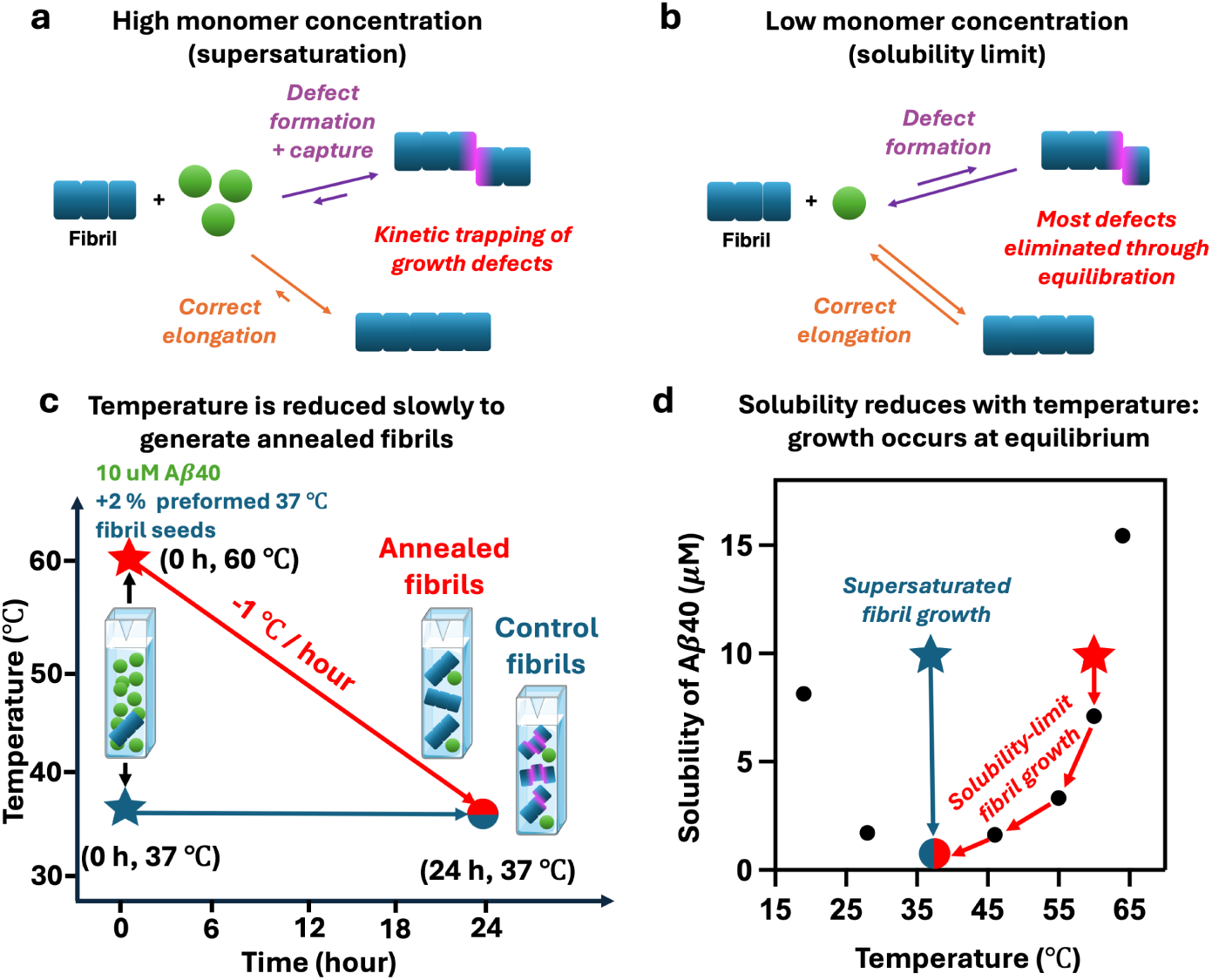
Controlling fibril growth defect frequency by varying supersaturation levels. **a**: defects are initially formed by incorrect or incomplete binding of an incoming monomer onto the fibril end and subsequently kinetically trapped by attachment of further monomer layers on top of the misaligned monomer. Defects have been illustrated as lateral dislocations of the entire cross-section for convenience only: other defect structures are also possible. **b**: at low enough monomer concentrations the rate of correct elongation and dissociation are close to equilibrium (the solubility limit). In this regime defect removal is faster than formation, and kinetic trapping is largely eliminated. **c**: we grow fibrils designed to be largely free of growth defects, by slowly reducing the temperature of a solution of initially slightly supersaturated monomeric Aβ40 from 60 to 37℃ by 1℃ per hour. **d**: the accompanying gradual decrease in solubility with temperature ensures elongation remains close to equilibrium at all times. The black data points are adapted from [39].

Defect stoichiometry can therefore in principle be reduced by eliminating their kinetic trapping. Although equilibrated solutions of mature fibrils are not kinetically trapped by high supersaturation, the prevalence of long fibrils (often containing *>* 10^5^ monomers) leads to very slow exchange of protein in the interiors of fibrils with the residual monomeric protein in solution [30, 31, 41]. Therefore, fibril growth defects will remain strongly kinetically trapped unless experimental conditions are chosen to almost completely disassemble the fibrils. Instead, a more practical approach is to prevent kinetic trapping in the first place by growing the fibrils at very low supersaturation (Fig. 3**b**).

To do so, we took advantage of the fact that fibril solubility - the monomer concentration at which elongation and dissociation rates equalize - can be altered by changes in temperature (Fig. 3**d**). We measured the solubility of Aβ40 fibrils at 60℃ in 20 mM sodium phosphate, pH 7.4 by NMR, finding it to be 7.1 µM (Fig. SI3). This solubility is much higher than the measured ca. 400 nM at 37℃ [42] and is in agreement with a previous study that found a steady increase in Aβ40 solubility with temperature above 37℃ [39]. A gradual reduction in temperature from 60 to 37℃ of a monomeric solution at a concentration near the 60℃ solubility value should then enable fibril growth at almost zero supersaturation (Fig. 3**c**). Such *annealed* fibrils are expected to have far fewer growth defects than fibrils produced from a similar (and therefore highly supersaturated) initial monomer concentration at 37℃. This approach was inspired by the common manufacturing technique of *annealing* which removes defects in metals formed during forging or in self-assembled polymer and lipid bilayer systems [43–45]. Here, we worked with Aβ40 rather than Aβ42 because Aβ40 has a much higher solubility, making it easier to generate sufficient annealed fibril concentrations at low supersaturation.

We generated a first generation of Aβ40 fibrils by allowing a solution of 57 µM Aβ40 monomer to fully aggregate at 37℃ (Fig. SI4). Second-generation annealed and control fibrils were produced by incubating 0.2 µM monomer-equivalent of the first generation fibrils with 10 µM fresh Aβ40 monomer. To make annealed fibrils, the incubation temperature was started at 60℃ and was reduced by 1℃ every hour until reaching 37℃ (Fig. 3**c** and SI Sec. 4c). To make control fibrils, the incubation temperature was maintained at 37℃ throughout. Seeding bypasses primary nucleation [35]. Moreover, at the low supersaturation levels used to make annealed fibrils, secondary nucleation becomes much less important than elongation. Therefore, most annealed fibrils are expected to be elongated first generation seeds. Since elongation typically preserves fibril morphology [46], using first generation seeds therefore greatly reduces the risk that the annealing procedure leads to the formation of fibrils with different morphology compared to those formed at 37℃.

To confirm that the annealing procedure does not cause morphological changes, we performed high-resolution cryo-electron microscopy on both annealed and control fibrils. We observed no significant differences in the raw images (Fig. 4**a**-**b**). We then performed 2D class averaging and once again there are no detectable differences between the collated 2D class averages (Fig. 4**c**-**f** ). We finally collected statistics on these classes, and found that the proportions of twisted vs straight fibrils were almost identical in each sample (43:57 in annealed fibrils and 40:60 in control fibrils). Furthermore, the average crossover length was *>* 300 nm in both samples, and the fibril widths were identical to within error. In other words, there are no detectable morphological differences between annealed and control fibrils. The only possible major differences between the annealed and control fibrils are therefore fibril length, and defect frequency and structure.

**Fig. 4.**
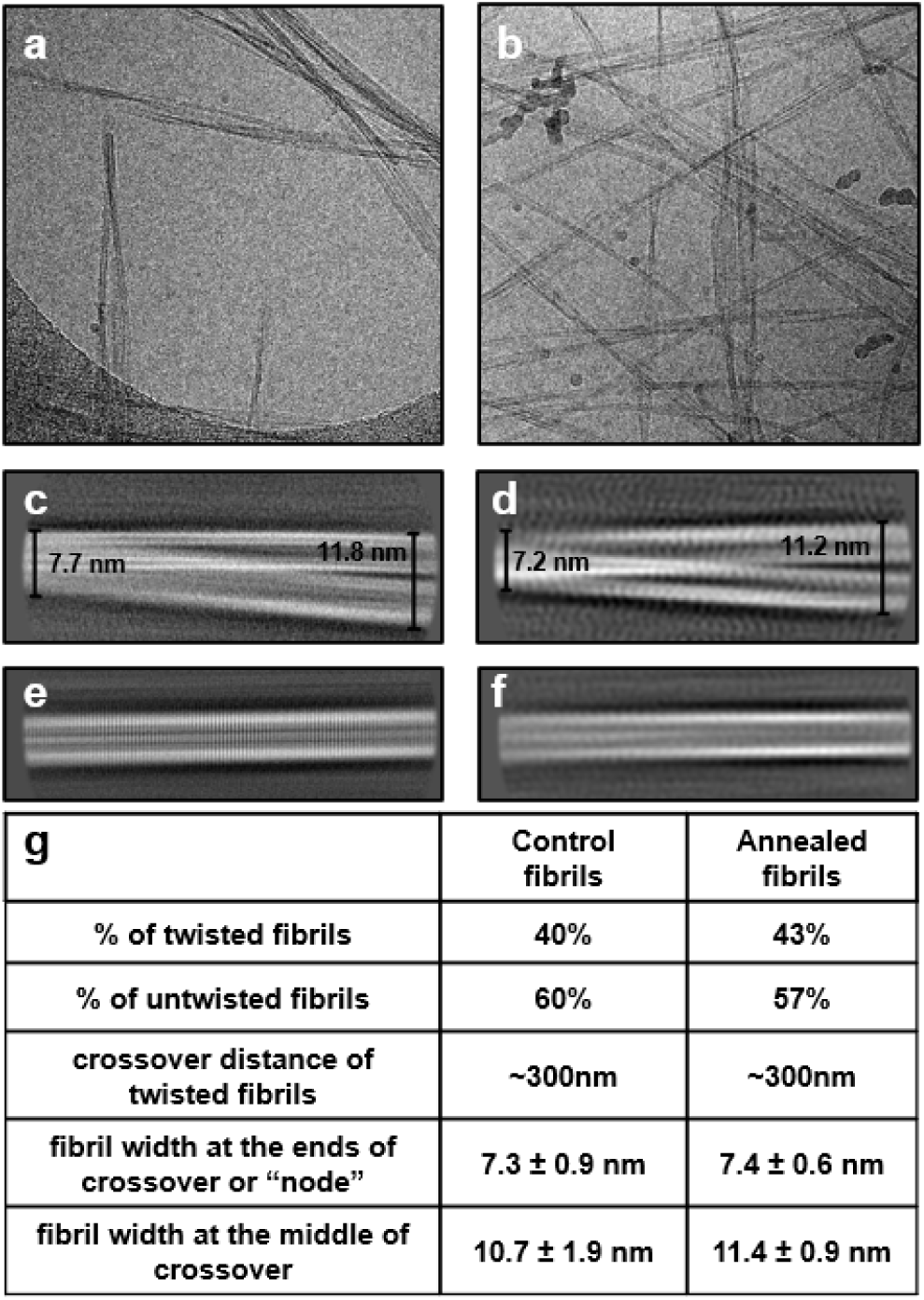
Cryo-EM data processing and statistics of control fibrils vs annealed fibrils. **a** and **b**: motion corrected micrographs showing control fibrils and annealed fibrils, respectively. **c** and **d**: 2D class averages for twisted fibrils in control fibrils and annealed fibrils datasets, respectively, and measurements of fibril diameter at points along the fibril crossover. **e** and **f** : 2D class averages for untwisted fibrils in 37℃ fibrils and annealed fibrils datasets, respectively. **g**: summary of fibril morph population and morphology statistics.

It must be emphasized that high-resolution cryo-electron microscopy algorithms by design can only reconstruct regular, periodic structural features of amyloid fibrils. Indeed, it is precisely the predominant regularity and periodicity of amyloid fibrils that allows such algorithms to function and to reconstruct high-resolution structures. Rare, irregular defects can never be visualized using current software no matter how large a sample is analyzed, as irregularities appear only as noise to the algorithms and are therefore averaged away during image processing [47]. Therefore, although cryo-EM can rule out morphological differences between our samples, it can never directly provide information about numbers and structures of defects. Note that, due to current software limitations, the average crossover length is too large for 3D fibril structures to be calculated; however, in this study these 3D structures are not needed since the 2D classes already rule out significant structural differences in the fibrils.

Growth defects are also expected to be too small (occupying 1-2 fibril planes) to be directly visible in raw cryo-EM images prior to application of high-resolution structure determination algorithms. However, certain types of growth defects can be identified indirectly by their nonlocal effects on fibrils. These include branching sites, and defects that punctuate transitions from one fibril morphology to another possessing a different twist length, both of which are clearly identifiable in raw cryo-EM images of Aβ40 and Aβ42 fibrils (see SI Figure S5). Other types of growth defects such as ordinary dislocations affect only the directly adjacent planes of the fibrils and are therefore not expected to be visible even indirectly in these samples. Any of these defects, whether visible or invisible, could plausibly act as secondary nucleation sites.

### 2.4 Annealed fibrils possess far fewer secondary nucleation sites than normal fibrils

Since Brichos also strongly inhibits Aβ40 aggregation at very low stoichiometries [48], it is likely that, as was shown for Aβ42 fibrils, Brichos binds almost exclusively to secondary nucleation sites and not elsewhere along Aβ40 fibrils. We therefore again sought to quantify the number of secondary nucleation sites for both annealed and control fibrils by measuring the Brichos binding stoichiometry to Aβ40 fibrils. This time we used fluorescence correlation spectroscopy (FCS) instead of MDS to eliminate the possibility that low recorded Brichos binding stoichiometries are an artifact of either experimental technique. We mixed different concentrations of Alexa488-Brichos with fixed concentration of non-fluorescent Aβ40 fibrils. We measured free Alexa488-Brichos concentration by positioning the confocal volume at a height where only diffusible Alexa488-Brichos can be detected, excluding those attached to the sedimented fibrils at the bottom of the dish (Fig. 5**a**). Visual inspection of the resultant binding curves of free vs total Brichos (Fig. 5**a**) reveal annealed seeds to have a lower Brichos saturation concentration compared to control seeds, indicating a lower secondary nucleation site frequency, and supporting the hypothesis that they are growth defects.

**Fig. 5.**
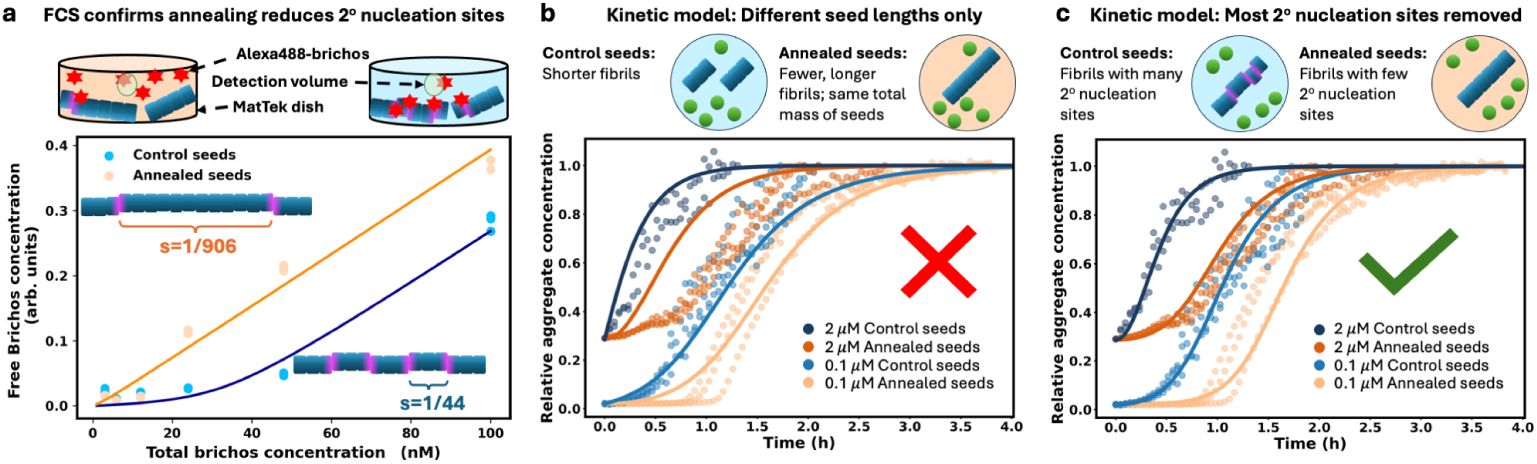
Annealed fibrils have fewer secondary nucleation sites and lower secondary nucleation rate than control fibrils. **a**: Solutions of 4.5 µM Aβ40 fibrils (annealed or control) with 3 to 100 nM Alexa488-Brichos are incubated for 2 h at 37*^◦^*C, and monitored by FCS to see the number of diffusing Alexa488-Brichos. Fitting a standard ligand-protein 1 : 1 binding model (Eq. (1)) reveals the number of Aβ40 molecules in the fibril per Alexa488-Brichos binding site for annealed seeds and for control seeds is *s*=1/44 (R² = 0.969) and *s*=1/906 (R² = 0.974), respectively. **b** and **c**: The aggregation of 5 µM Aβ with 0.1 or 2 µM monomer-equivalent of annealed or control seeds was monitored by ThT fluorescence in a plate reader. The fitting (lines) assumed that the delayed aggregation in the presence of annealed seeds compared to control seeds is due either to longer fibril length (**b**) or to fewer secondary nucleation sites (**c**) in annealed seeds compared to control seeds (see SI Sec. 2 for kinetic models). Again, defects have been illustrated schematically as lateral dislocations for convenience only.

We then fit these binding curve data using a simple protein-ligand binding model [49](Eq. (1)). We found one Brichos binding site per 44 monomers in control fibrils. This, alongside the fact that an entirely different site quantification technique was used compared to Aβ42, provides even more support for the rarity of Aβ secondary nucleation sites. In annealed fibrils, the stoichiometry of these sites was reduced to one per 906 monomers. This implies that the number of secondary nucleation sites are reduced by approximately 95% by the annealing treatment relative to fibrils formed entirely at 37℃. This large reduction validates a key prediction of the hypothesis that the secondary nucleation sites are growth defects.

Note that, if Brichos also binds elsewhere along Aβ40 fibrils, the secondary nucleation site stoichiometry would be even lower than what was inferred here from the Brichos binding stoichiometry. Since the annealed and control fibril morphologies have been demonstrated to be the same on average, this would in turn imply an even greater reduction in secondary nucleation site incidence in annealed fibrils compared to control fibrils if they are indeed growth defects.

### 2.5 Annealed fibrils display considerably reduced secondary nucleation seeding efficiency compared to normal fibrils

We next used 0.1 or 2 µM monomer-equivalent of the second generation fibrils (control seeds or annealed seeds) as preformed seeds in new aggregation reactions of 5 µM monomeric Aβ40 at 37℃. We find that even though both control and annealed seeds accelerate the aggregation (Fig. SI1), the annealed seeds have much less seeding efficiency compared to control seeds in both lightly and heavily seeded conditions, Fig. 5 **b**-**c**.

The decrease of seeding efficiency of annealed seeds compared to control seeds could theoretically be due to the annealed seed fibrils having greater length, and therefore fewer fibril ends where elongation can occur for an equivalent fibril mass. Producing the second generation seeds by seeding with first generation fibrils will reduce but not eliminate these length differences by reducing the role of nucleation relative to elongation in their formation kinetics. To test this possibility we therefore first attempted to fit the data using the standard kinetic model for Aβ40 aggregation [14], but where the seed fibril length was allowed to differ between control seeds and annealed seeds (i.e., we allowed the seed fibril number concentration *P*_0_ to vary, but not the seed mass concentration *M*_0_). This model was unable to globally fit the data, even with extreme (≫ million-fold) length differences, yielding misfits in particular to the kinetic curves for the reactions with 2 µM monomer-equivalent of preformed fibril seeds (Fig. 5 **b**). Therefore, the possibility that control seeds are shorter compared to annealed seeds due to fragmentation is excluded. (Note that this could not be tested by sonicating the seeds to the same length, because it is likely that fibrils might preferentially break at defects, in which case sonication would affect defect stoichiometry as well as fibril length. Moreover, Aβ40 fibrils are resistant to sonication [50].)

The reduced number of secondary nucleation sites in annealed fibrils can be captured by a kinetic model that is identical in all respects to the standard model for Aβ40 aggregation [14], but where the seed mass concentration *M*_0_ is multiplied by fitting parameter *r*, the fraction of remaining secondary nucleation sites compared to control fibrils (See SI Sec. 2 for modified kinetic model equations). We find that such a model, where effectively *M*_0_ can vary but not *P*_0_, can fit the measured kinetic data very well (Fig. 5 **c**), using a value for *r* of 0.08, which is consistent with a 92% reduction in secondary nucleation site numbers. This is remarkably consistent with the 95% reduction estimated using Brichos stoichiometry and is despite this model having no more degrees of freedom than the differing-seed-length model. The close correspondence between secondary nucleation site stoichiometry and secondary nucleation propensity implies that there is little if any structural difference between the secondary nucleation sites in annealed and control fibrils.

Brichos binding can increase the half-time of Aβ40 aggregation by at least 10- fold [48]. Because the half-time depends on the inverse square root of the secondary nucleation rate [51], this indicates higher concentrations of Brichos can reduce secondary nucleation by at least 100-fold, and possibly much more. The maximum possible contribution of non-defect sites that are not bound by Brichos to the overall secondary nucleation rate is therefore 1%. We have found that Aβ40 fibrils formed without supersaturation have approximately 5% of the defect sites possessed by fibrils grown at typical supersaturation levels. Secondary nucleation on defects is therefore still at least 5 times faster than secondary nucleation from other non-defect sites even in annealed fibrils. So, fibril growth defects should remain the dominant source of Aβ40 secondary nucleation regardless of supersaturation levels.

### 2.6 Fibril growth defect stoichiometry and stability

At equilibrium, the fraction of defective fibril planes versus total fibril planes *p_eq_* is mathematically related to the free energy penalty Δ*G*_def_ for forming a dislocation defect by −*RT* ln(*p_eq_*) = Δ*G*_def_ (assuming defective layers are rare and predominantly of only one structure, see SI Sec. 1). *p_eq_* is related to the equilibrium defect site stoichiometry per monomer *s_eq_* by *s_eq_* = *n*_def_ *p_eq_/x*, where *x* is the number of monomers in the fibril cross-section and *n*_def_ the number of discrete defect sites per defective layer (SI Sec. 1). For instance, *n*_def_ = 2 in dislocation-type defects because dislocations expose fibril core on both sides of the fibril.

The average annealed Aβ40 formation temperature is approximately the temperature at which half of the monomer has been aggregated. If protein aggregation were instantaneous, the solubility vs temperature chart (Fig. 3**d**) would imply this to be around 57℃. Accounting for the finite time taken for protein aggregation starting from a supersaturated solution, it is probably closer to 55℃. The defect stoichiometry in annealed fibrils, *s* = 1*/*906, is therefore approximately *s_eq_* at 55℃. In SI Sec. S1 we use these numbers to estimate the equilibrium defect free energy penalty for several plausible values of *x* and *n*_def_, finding Δ*G*_def_ to range from -14.8 to -18.6 kJ/mol. This being many multiples of the thermal fluctuation energy *RT* , these defects are thus very thermodynamically unstable, as expected given their low stoichiometry. Δ*G*_def_ is also proportionally significant: approximately half of the per-monomer free energy for correct elongation (calculated in SI Sec. S1 as -34.4 kJ/mol at 55℃).

We can also predict that proteins with higher solubilities will have a higher defect incidence in their fibrils because weaker intermolecular interactions in the fibril will correspond to a smaller free energy penalty for a growth defect. Our finding that the higher solubility Aβ40 has higher defect incidence in its fibrils compared to Aβ42 is consistent with this prediction. For instance, α-synuclein has a solubility of 4.1 µM at the conditions used in ref [52] (37℃ and pH 7.4), corresponding to a per-monomer free energy of fibril elongation of -32 kJ/mol, compared to -38 kJ/mol for Aβ40 under these conditions. A dislocation defect exposing an equal number of monomers, or a partial misfolding defect with a similar number of amino acids involved, will therefore likely incur a lower free energy penalty in α-synuclein fibrils, and consequently appear at a higher frequency.

## 3 Discussion

There are several reasons to believe that defects may act as secondary nucleation sites across many or even most types of amyloid fibrils. There exists a variety of amyloid-forming proteins, and the surfaces of the diverse fibrils they form are composed of a similar variety of peptide sequences. About the only structural feature these fibrils have in common is their hydrogen bonding between adjacent planes in the filaments, with significant differences in the packing of hydrophobic side-chains in the fibril core. The universal nature of secondary nucleation itself across amyloid-forming proteins [9, 53], independent of fibril surface structure and properties, thus already suggests that secondary nucleation may have more to do with the amyloid core than with the fibril surface. Indeed, it has not yet been found to be possible to design an Aβ42 mutant that eliminates secondary nucleation without also eliminating fibril growth [16]. Moreover, Aβ42 mutations at residues not involved in the amyloid fibril core do not strongly affect the ability of fibrils composed of these mutants to catalyze secondary nucleation of WT Aβ42 [54, 55]. Another structural feature shared by all amyloid fibrils is the presence of growth defects, which is guaranteed by the laws of thermodynamics (the Boltzmann distribution) regardless of their exact structure. Since these defects can provide partial access to the fibril cross-section, they are a very strong candidate for secondary nucleation sites. Such defects can provide a partial scaffold to surmount the entropic penalty associated with forming a new oligomer or fibril [56–58]. Additional support for this proposal is found beyond the field of amyloid fibril self-assembly: secondary nucleation in metal and polymer crystallisation has also been observed to occur at defects generated during crystal growth [59–61], whose formation can be similarly promoted by increasing supersaturation.

A recent study [52] found Brichos to strongly inhibit secondary nucleation at pH 7.4 at a molar ratio of Brichos:α-synuclein of 1:10. Similarly, Brichos strongly inhibits IAPP by binding secondary nucleation sites [62]. The observed *>* 10-fold increase in aggregation half time implies a reduction in secondary nucleation rate of over 99% at a molar ratio of Brichos:IAPP of 1:10 (since half-times scale as the inverse square root of the secondary nucleation rate constant [51, 63]). The stoichiometry of secondary nucleation sites is therefore at most 1/10 in both cases, and are likely structural defects, similarly to Aβ40 and Aβ42 fibrils. The ability of Brichos to bind tightly to secondary nucleation sites in Aβ40, Aβ42, α-synuclein and IAPP fibrils implies these sites must have common structural features. Given how unrelated these proteins are and thus how unrelated the surfaces of their amyloid fibrils must be, this commonality could be that they are all defects that partially expose cross-β core structure. If Brichos indeed has generic binding affinity for cross-β structure, and if growth defects universally promote secondary nucleation, then it could be a truly universal inhibitor of secondary nucleation. This universal affinity for cross-β structure would also explain why Brichos often displays at least some ability to inhibit elongation by binding fibril ends [52, 64], which should have similar (although not identical) structure to the defect sites.

Typically, surfaces catalyze the nucleation of amyloid fibrils and nucleation more generally. Indeed, primary nucleation of fibrils (and unrelated aggregates) almost always happens on interfaces *in vitro*, not in bulk [53, 65, 66]. It stands to reason that other areas of the fibril surface can also act as secondary nucleation sites, in addition to growth defects. Such “off-defect” secondary nucleation, and its inhibition by Brichos, has recently been proposed as the dominant mechanism for secondary nucleation of α-synuclein at pH 4.8 [67], although the experimental evidence presented is also consistent with defect-driven nucleation. Note this surface-diffusion-based off-defect mechanism is certainly ruled out for Aβ42 and Aβ40 secondary nucleation by the much lower stoichiometry and *K_D_* of Brichos binding to Aβ fibrils compared to α-synuclein fibrils at pH 4.8, as well as the detailed inhibition kinetics as shown in [23]. Regardless of whether significant off-defect nucleation occurs, we expect that both monomeric amyloid proteins and Brichos can bind to both defect sites and other areas of the fibril surface, albeit likely with very different affinities. Indeed, in the case of Aβ42 fibrils, two distinct binding sites for Brichos have recently been demonstrated [27], one tight (with a low dissociation constant *K_D_* of 12.9 nM at 37℃) and one loose (*K_D_* = 18.5 µM at 37℃). The dissociation constant for Brichos on Aβ42 secondary nucleation sites at 37℃has previously been calculated as 40 nM [68], implying only the tight binding sites are important for secondary nucleation. Similarly, two binding sites for Bri2- Brichos on α-synuclein fibrils at pH 7.4 have recently been demonstrated [52], one tight (*K_D_* = 22 nM at 25℃) and one loose (*K_D_* = 350 µM at 25℃). When only one binding site is assumed in the modelling of the binding experiments, an average *K_D_* value of 2.3 µM is recovered in [52], similar to the *K_D_* values reported in [67] for different ProSP-C Brichos variants at pH 4.8 (0.45–1.3 µM at 21℃). Also, a loose binding site for α-synuclein monomers on α-synuclein fibrils of *K_d_*∼ 1 mM has been reported in [69] at pH 7 (temperature not stated). This can be contrasted with a *K_D_ <* 10 µM for α-synuclein monomers specifically on secondary nucleation sites at pH 5.5, that can be inferred from the complete saturation of secondary nucleation at 37℃above a monomer concentration of 10 µM [70]. Although not directly comparable due to the different pH’s and consequent probable different fibril morphologies, the *>* 100-fold difference in *K_D_* is nonetheless suggestive of different types of binding sites. (Note Aβ40 and Aβ42 monomers have similar *K_D_* values for Aβ40 and Aβ42 fibril secondary nucleation sites of around 6 µM.)

Crucially, the finding that secondary nucleation sites are defects could unlock a new approach for developing Alzheimer’s and Parkinson’s disease therapeutics using structure-based drug design. Secondary nucleation sites on fibrils have long been recognized as a promising target for rational design of therapeutics [24], being both the major source of toxic amyloid-β oligomers [71], and obligate for rapid fibril proliferation [9]. Indeed, much of the interest in Brichos itself stems from its ability to bind to and block these sites to reduce the rate of secondary nucleation [72]. It also implies that targeting supersaturation levels in the brain (by e.g. reducing amyloidogenic protein production) will be an even more effective alternative therapeutic approach than anticipated, since it will not only directly lower aggregation rates but also result in fewer defects in the formed fibrils, which in turn could lead to the formation of fewer toxic oligomers.

It will be important to investigate the relationship between fibril morphologies and growth defects in future. The fibril core structure exposed at a defect could potentially be templated onto the newly forming fibril during secondary nucleation, offering a possible molecular pathway for strain propagation. A recent investigation of Brichos binding to Aβ42 fibrils [27] found that full inhibition of secondary nucleation by Brichos gives rise to a different, thinner morphology with a single filament consisting of two monomers per plane, as opposed to the more usual two-filament morph. This could indicate that secondary nucleation of Aβ42 propagates structure in some way. Alternatively, it could point to a key role of growth defects such as dislocations in anchoring the higher-order assembly of filaments. Additionally, it is plausible that growth defects in other types of fibrils enable fibril fragmentation instead of or as well as secondary nucleation, since the bonding is necessarily weaker at defect sites. If so, and considering that filament branching is also a defect-enabled phenomenon, the formation of growth defects could be the key molecular event driving autocatalytic amplification in all types of fibril formation reactions.

As explained above, if fibrils are grown using initial monomer concentrations well above the solubility, as is usual in kinetic experiments, growth defect incidence will often be much higher than that predicted by the Boltzmann distribution. It must be emphasized that these excess defects are nonetheless kinetically very stable even after supersaturation has been removed and therefore persist in mature fibrils for very long periods of time. This extremely strong kinetic trapping of excess defects arises from the great length of fibrils, with the average defect many thousands of monomers distant from the fibril end. The need to depolymerise tens of thousands of monomers from the fibril to access the average defect imposes an insurmountably enormous kinetic free energy barrier for defect removal. Although defects are transiently near fibril ends during their initial formation, when fibrils are assembled under highly supersaturating conditions new monomers add on top of defective fibril layers far faster than defective fibril layers can disassemble and re-assemble in the correct structure. Experimental confirmation of the slow exchange of monomers in the interior of fibrils with those in solution (the cause of the strong kinetic trapping) is abundant [73, 74].

In summary, our study shows that amyloid-β fibrils contain a small number of secondary nucleation sites, which can be further reduced by decreasing the level of supersaturation. These nucleation sites are identified as defects formed during fibril growth. The free energy penalty for a growth defect is roughly 50% of the per-monomer elongation free energy. A deeper understanding of defect-driven secondary nucleation could open new opportunities for targeting amyloid formation in various diseases. By inhibiting secondary nucleation or reducing fibril defects, it may be possible to lower the formation of toxic species and slow disease progression. These findings offer a broader perspective on the molecular mechanisms governing amyloid-related diseases and offers a potential starting point for structure-based drug design for inhibiting amyloid secondary nucleation.

## 4 Methods

### Chemicals and consumables

All Aβ42 or Aβ40 related experiments were carried out in 20 mM sodium phosphate buffer (sodium dihydrogen phosphate dihydrate and disodium hydrogen phosphate) at pH 8.0 or pH 7.4, respectively. The buffer components were purchased from Sigma-Aldrich (St. Louis, MO, USA), and all buffer solutions were prepared using Milli-Q water. The buffer was filtered through wwPTFE 0.2 µm 50 mm disc filters (Fisher Scientific, Pittsburgh, PA, USA) and degassed prior to use. The dye Thioflavin T (ThT) for monitoring amyloid formation was obtained from Calbiochem. Axygen™ MaxyClear Snaplock Microtubes and Corning® 96-well Half Area Black/Clear Flat Bottom PEGylated Polystyrene Microplates (3881) were used as containers for protein solutions to minimize protein adsorption onto container surfaces.

### Expression and Purification of Aβ42, Aβ40 and Brichos

The expression and purification procedure of the proSP-C Brichos domain, the peptide Aβ(M1- 42) (MDAEFRHDSGYEVHHQKLVFFAEDVGSNKGAIIGLMVGGVVIA), referred to here as Aβ42 and A*β*(1-40), referred to here as Aβ40 (DAEFRHDSGYEVHHQKLVFFAEDVGSNKGAIIGLMVGGVV) in *E. coli* have been described previously [48, 75, 76]. The purified Aβ42 or Aβ40 powder was stored at -20*^◦^*C.

Immediately prior to each experiment, Aβ42 or Aβ40 powder was dissolved in 6 M guanidine hydrochloride (pH 8.5), and monomers were isolated by size exclusion chromatography (Superdex 75 column) in 20 mM sodium phosphate buffer as described in chemicals and consumables session.

ProSP-C Brichos domain was covalently labelled with the Alexa-488 dye (Thermo Fisher Scientific, Waltham, US) by incubating the protein for three hours in 20 mM sodium phosphate buffer, pH 8.0, at room temperature under gentle agitation with one equivalent of Alexa-488 carboxylic acid succinimidyl ester to label covalently primary amines. Following the labelling reaction, excess dye was removed by desalting on a G25 gel filtration column, followed by purification by anion exchange to remove any protein with multiple labels, and an additional desalting step on a G25 gel filtration column before use.

### Microfluidic Diffusional Sizing

The fabrication and operation of the microfluidic diffusion device were performed as described previously [32, 33]. Briefly, the microfluidic chips were fabricated by using standard soft lithography techniques into polydimethylsiloxane [77]. The analyte sample and the buffer were introduced in the device through reservoirs connected to the inlets. The flow rates in the channels were controlled by applying a negative pressure at the single outlet channel by means of a syringe pump (neMESYS, Cetoni GmbH, Korbussen, Germany). Typical flow rates were in the range 60 µl/h to 90 µl/h and lateral diffusion profiles were recorded at twelve different positions (3.52 mm, 5.29 mm, 8.57 mm, 10.33 mm, 18.61 mm, 20.37 mm, 28.65 mm, 30.41 mm, 58.69 mm, 60.45 mm, 88.73 mm and 90.5 mm) by standard epifluorescence microscopy using a cooled CCD camera (Evolve 512, Photometrics, Tucson, AZ, USA). The diffusion profiles were fitted by model simulations based on advection-diffusion equations under the assumption that the diffusivities of the species in the mixture are distributed according to a bimodal Gaussian distribution [32]. In this approach, the fitting parameters are the mean and the standard deviation of at least two size distributions, corresponding to the populations of the free chaperone (small size range) and of the chaperone bound to the fibrils (large size range).

In order to evaluate the binding between Brichos and Aβ42 fibrils during the aggregation process, samples with Alexa-488 labeled Brichos at concentrations of 0.25, 0.35 or 0.50 µM were mixed with 24 µM Aβ42-monomers and 20 µM ThT and incubated in a 96-well plate in a Fluostar Optima plate reader (BMG Labtech, Ortenberg, Germany) at 37*^◦^*C under quiescent conditions. The aggregation kinetics were monitored by following the associated increase in ThT fluorescence. After completion of the reaction, the samples were incubated for at least 2 days at 21*^◦^*C to ensure equilibrium conditions. They were subsequently analysed by microfluidic diffusion sizing at 21*^◦^*C. To evaluate the binding between Brichos and mature Aβ42 fibrils, the same Aβ42 aggregation reaction was performed at 37℃ but without Brichos. After completion of the reaction, aliquots of Alexa-488 labeled Brichos in the concentration range between 0 and 1.5 µM were added to samples of the mature Aβ42 fibrils and incubated for at least 2 days at 21*^◦^*C to ensure equilibrium conditions. They were subsequently analysed by microfluidic diffusion sizing at 21*^◦^*C.

### ThT kinetics

The kinetic assay for Aβ40 displayed in Fig. 5**b**-**c** starts with 5 µM monomer in 20 mM pH 7.4 phosphate buffer with 10 µM ThT, with or without 0.1 or 2 µM monomer-equivalent of seeds. The seeds were generated as described in the static light scattering section (SI Sec. 6). After mixing the monomer and the seeds, the solutions were immediately incubated in a 96-well half-area clear flat bottom PEGylated polystyrene microplate at 37*^◦^*C. The fluorescence was recorded in a Fluostar Omega plate reader (BMG Labtech) using a 440 nm excitation filter and a 480 nm emission filter. The waiting time between the reading cycles was set to 0, meaning that the wells were continuously monitored throughout the kinetic measurements. This setup is designed to control the agitation frequency caused by reading wells consistently across all experiments [38].

The kinetic assays for Aβ42 were performed by incubating a solution of 3 µM Aβ42 peptide, 1% seeds, and 20 µM ThT with or without different concentrations of Brichos in a non-binding 96 well plate in a plate reader Fluostar Optima (BMG Labtech, Ortenberg, Germany) at 37*^◦^*C with double orbital rotation (400 rpm).

### Fluorescence Correlation Spectroscopy

For FCS measurements, 10 µL samples of 4500 nM Aβ40 fibrils and 3-100 nM Alexa-488 Brichos were placed in MatTek dishes (35 mm, 10 mm glass bottom, No. 1.5 glass) at 37*^◦^*C . The measurements were conducted using a Zeiss 780 confocal laser scanning microscope equipped with a Zeiss C-Apochromat 40×/1.2 NA water immersion objective. Excitation was performed at 488 nm, and fluorescence emission was collected within the 499–622 nm range. The free Brichos concentration was determined by fitting the correlation curves at a time range of 10*^−^*^5^ to 1 s to theoretical models for diffusion of two species of different molecular weights, representing the diffusion of free dye and Brichos(Fig. SI2) [78]. Each data point in Fig. 5 **a** is based on 3-4 FCS measurements, 90 s measurements each.

To determine the number of secondary nucleation sites per monomeric residue (*s*), the free Brichos concentration *y* as a function of total Brichos concentration *x* was fitted with the following ligand binding equation, derived in the SI:

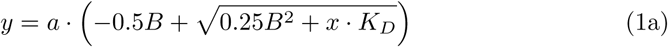

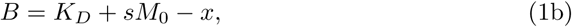

where *a* is an arbitrary proportionality constant, *K_D_* is the dissociation constant for Brichos binding, and the quantity in the brackets is the concentration of free Brichos. The concentration of Aβ40 fibrils is *M*_0_ = 4500 nM. We globally fitted *K_D_* obtaining 2.3 nM, and *a* = 0.004; the other parameters were allowed to differ between control and annealed fibrils. We obtained *s*= 1*/*44 for control fibrils and *s*= 1*/*906 for annealed fibrils.

### Cryo-EM sample preparation

The samples were imaged using cryo-EM as described previously [76]. Specimens were plunged using an automated plunge freezer (Leica EM GP) set at 20 *^◦^*C with 90% relative humidity. Thin liquid films were created on glow-discharged lacey carbon 300 mesh grids (Agar Scientific) and blotted with filter paper before being plunged into liquid ethane at -184*^◦^*C, ensuring vitrification in a glass-like state that prevents ice crystal formation and preserves the original microstructures. The specimens were stored in liquid nitrogen until further use.

### Cryo-EM imaging

The prepared specimen grids were transferred into the electron microscope (JEM 2200FS) using a Fischione model 2550 cryo-transfer tomography holder. Imaging was performed at an acceleration voltage of 200 kV, with zero-loss images captured digitally using a TVIPS F416 camera and SerialEM under low-dose conditions with a 10 eV energy filter. Image contrast was adjusted automatically using ImageJ.

### Cryo-EM data collection and processing

The cryoEM datasets were collected at the SciLife lab node in Umeå, Sweden, using a Titan Krios electron microscope (Thermo Fisher) operated at 300 kV with a Falcon4 detector and a Selectris energy filter (Thermo Fisher) was used operating with a 10 e*^−^*V slit. Datasets of size 8106 movies and 7990 movies were collected for control and annealed fibrils respectively, at a nominal magnification of 130,000x was set yielding a pixel size of 0.92 Å. A defocus range of -1.4 to -2.6 µm and a total dose of 40 e*^−^*/Å^2^ were used over an exposure time of 4.39 seconds.

The raw EER movies were fractionated, aligned and summed using motion correction in RELION-4 [79] and CTF estimation was done for micrographs using CTFFIND4 [80]. Manual picking of fibrils was done until roughly 100,000 particles of 300 pixel box size were picked, and the extracted segment was used to train a separate picking model for each dataset in Topaz, which is incorporated in RELION-4.0 [79]. Iterative rounds of 2D classification were performed to remove picking artefacts such as carbon edges and ice contamination, until only classes that could be protein fibrils remained. For 2D classification, a box size of 600 pixels was used. Twisted and straight fibrils were then separated out and a final round of 2D classification was performed separately for both sets of fibrils.

## Supplementary information

This article has an associated Supplementary Information file.

## Acknowledgements

This work was supported by the Swedish Research Council (2019-02397 to ES, 2015-00143 to SL), and the GenerationNano project, the European Union’s Horizon 2020 research and innovation program under the Marie Skl-odowskaCurie grant agreement No 945378 (S.L. co-PI). We acknowledge support from the Wellcome Trust (TPJK), the Cambridge Centre for Misfolding Diseases (TPJK), the BBSRC (TPJK), the Frances and Augustus Newman foundation (TPJK), the ERC PhysProt (agreement n 337969) (TS, TPJK, SL), ETC StG ”NEPA” (AS), the Royal Society (SC, AS), the ERASMUS Programme (TS), and The Danish Council for Independent Research — Natural Sciences (FNU-11-113326) (MA). This work was also funded by the Novo Nordisk Foundation (#NNF19OC0054635 to S.L.). We are grateful to the late Professor Sir Christopher Dobson for invaluable conversations regarding the microfluidic diffusional sizing experiments. We are also grateful to Quentin A. E. Peter and Thomas Müller for their guidance on microfluidic device design.

