## Supporting Information for "Structural defects in amyloid-β fibrils drive secondary nucleation"

### 1 Equilibrium thermodynamics of fibril growth defects

Amyloid fibrils consist of stacked layers of small numbers of monomers, typically 2-5, typically with tens of thousands or millions of layers per fibril. Given the periodic symmetry along the long axis of (defect-free) fibrils, it is easiest to view the thermodynamics of their assembly layer-by-layer. Instead of considering discrete defect sites, we therefore consider defective layers, which may come in a variety of forms. They could contain one or more partially misfolded monomers (misfolded from the perspective of the regular amyloid structure). They could also be offset or misaligned compared to the preceding layer, but contain otherwise largely correct bonding within the layer. However, we do not need to make any assumptions as to the precise structure of the defective layer, other than that it is not so irregular as to prevent subsequent layers of monomers binding in the correct conformation for the regular structure. For the defect to become kinetically trapped, and located far from the fibril ends, this is in fact a requirement rather than an assumption.

#### 1.1 Derivation of equilibrium defect stoichiometry formulae

We denote the free energy of forming a new layer as $G_i$, where $i = 0$ denotes a layer of the regular structure, $i = 1$ is the most-stable-possible defective layer that allows subsequent regular growth, $i = 2$ is the next-most-stable-possible defective layer, etc. At equilibrium, the probability for including a given growth defect in a fibril is then:

$$p_{eq,i} = \frac{e^{-G_i/RT}}{\sum_j e^{-G_j/RT}}. \tag{1}$$

If a defective layer of type $i$ has $n_i$ discrete defect sites, then the overall defect site stoichiometry is:

$$s_{eq} = \frac{1}{x}\sum_i n_i \, p_{eq,i}, \tag{2}$$

where $x$ is the number of monomers in a layer. The sum can run from $i = 1$ or $i = 0$ since $n_0 = 0$.

Now, if defective layers are rare, which is equivalent to them being thermodynamically quite unstable compared to correctly-assembled layers (and quite likely *a priori*, given the highly regular nature of the amyloid structure), then the defect probability becomes:

$$p_{eq,i} \simeq e^{-\Delta G_i/RT}, \quad \Delta G_i = G_i - G_0. \tag{3}$$

Furthermore, if one type of defective layer is significantly more stable than others (or is a member of a class of defective layers of equal stoichiometry $n_i$ and similar $G_i$ values), then we can write:

$$p_{eq} = \sum_i p_{eq,i} \simeq p_{eq,1}, \tag{4}$$

and:

$$s_{eq} \simeq \frac{n_{\mathrm{def}}}{x} e^{-\Delta G_{\mathrm{def}}/RT}, \tag{5}$$

where we have for notational convenience defined $\Delta G_1 = \Delta G_{\mathrm{def}}$ and $n_1 = n_{\mathrm{def}}$. For instance, a lateral misalignment creates two defect sites, one on each side of the fibril. So, the equilibrium stoichiometry of defect sites per monomer is $s_{eq} = 2p_{eq}/x$.

Note, if the defective layer leads to weaker bonding both to the preceding and subsequently-assembled layers in the fibril, the bonding energy reduction in both planes must be included in $G_i$.

#### 1.2 Calculation of defect free energies

The measured defect site stoichiometry at equilibrium is $s_{eq} = 1/906$, and it is estimated that the average temperature at which these defect sites form is 55℃. This very low stoichiometry already implies that defects are unstable and that Eq. (3) therefore applies. Under the approximation that we ignore the contributions of all but the most stable type of defect, we can also use Eq. (5). The free energy penalty of formation of a defective layer is then

$$\Delta G_{\mathrm{def}} = RT \ln \left( s_{eq} \frac{x}{n_{\mathrm{def}}} \right). \tag{6}$$

We can now consider several cases.

##### 1.2.1 4-monomer-thick fibril

For a misalignment defect and a 4-monomer-thick fibril, $x = 4$ and $n_1 = 2$ and the free energy penalty per layer is:

$$\Delta G_{\mathrm{def}} = -16.7 \text{ kJ/mol}. \tag{7}$$

For a partially-misfolded single monomer defect, $n_1 = 1$ and the energy penalty is instead:

$$\Delta G_{\mathrm{def}} = -14.8 \text{ kJ/mol}. \tag{8}$$

For comparison it is useful to have a value for $G_0$. This can be calculated from the solubility $c_{eq}$, which at 55℃ is 3.32 µM [1]:

$$c_{eq} = K_D = e^{(G_0/x)/RT}. \tag{9}$$

Doing the calculation gives:

$$G_0 = -137 \text{ kJ/mol}. \tag{10}$$

note the bonding free energy for a single correctly-bound monomer is -34.4 kJ/mol at this temperature.

So, if the defect is a misalignment defect, it incurs approximately a 1/8 energy penalty, or approximately 50% of the bonding energy of a single correctly-bound monomer. If it is a single-monomer defect, it is slightly less than this at ca. 43%. Either way, these defective layers are clearly thermodynamically very unstable.

##### 1.2.2 2-monomer-thick fibril

For a misalignment defect and a 2-monomer-thick fibril, $x = 4$ and $n_1 = 2$ and the free energy penalty per layer is:

$$\Delta G_{\text{def}} = -18.6 \text{ kJ/mol.} \tag{11}$$

For a partially-misfolded single monomer defect, $n_1 = 1$ and the energy penalty is instead:

$$\Delta G_{\text{def}} = -16.7 \text{ kJ/mol.} \tag{12}$$

$G_0$ is now:

$$G_0 = -68.8 \text{ kJ/mol.} \tag{13}$$

These defective layer free energy penalties are slightly larger in absolute terms and much larger relative to the free energy for a correctly-bound layer.

Again, such defects are very thermodynamically unstable, with their free energy penalties being several multiples of $RT = 2.73$ kJ/mol.

#### 2 Kinetic model of A$\beta_{42}$ aggregation

We describe A$\beta_{42}$ aggregation using a chemical kinetics framework that accounts for the various microscopic steps of aggregation (Fig. 2(a) of main text), as described previously [2]. The unperturbed kinetic equations for the aggregate number concentration $P(t)$ and aggregate mass concentration $M(t)$ are

$$\frac{dP}{dt} = k_1 m^{n_1} + k_2\, m^{n_2} M\,, \tag{14a}$$
$$\frac{dM}{dt} = 2k_+ m\, P = -\frac{dm}{dt}\,, \tag{14b}$$

where $m(t)$ is the monomer concentration, $k_1$, $k_2$, $k_+$ are the rate constants for primary nucleation, secondary nucleation and elongation, and $n_1$, $n_2$ are the reaction orders of primary and secondary nucleation with respect to the monomer concentration.

Fits of the aggregation curves in the presence of Brichos were performed using the Amylofit platform (which implements analytical solutions to the kinetic equations (14a)-(14b)) [3], whereby the presence of Brichos was captured by means of an effective rate constant for secondary nucleation $k_2$ that depends on chaperone concentration.

When protein aggregation is seeded with annealed fibrils, Eqs. (14) still apply, but with modified initial condition $M(0) = rM_0$, where $M_0$ is the mass concentration of seed fibrils and $r$ is the fractional reduction in secondary nucleation sites caused by the annealing process. The analytical solution to such equations is then identical to that used in Amylofit [3], but with all instances of $M(0)$ replaced by $rM_0$. We used an offline version of Amylofit to fit this modified analytical solution to the data (without Brichos), and to determine $r$.

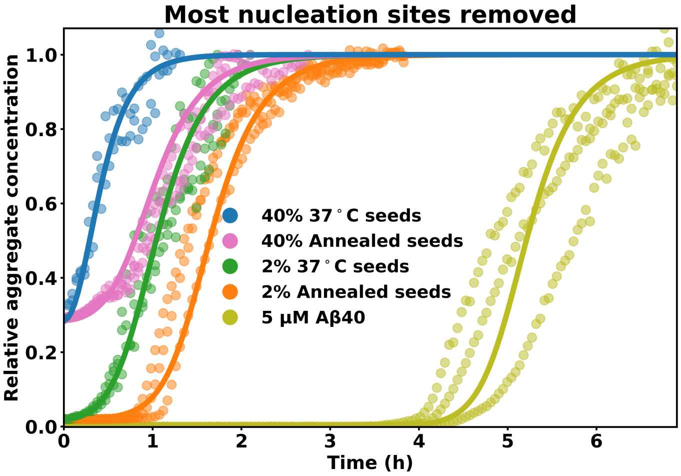


**Fig. 1** Aggregation of 5 $\mu$M Aβ40 at 37°Cwith 0%, 2% or 40% 37°C seeds or annealed seeds. The aggregation is monitored by ThT intensity. The fitting with the model of removing most nucleation sites is shown in lines.

### 3 FCS fitting

The fitting of the FCS data is shown in Figure 2**a** and **b**, using the model:

$$Y = \frac{1}{N}\left[\frac{1-f_2}{\sqrt{(1+\frac{X}{\tau_{D1}})(1+\frac{X}{\tau_{D1}S^2})}} + \frac{f_2}{\sqrt{(1+\frac{X}{\tau_{D2}})(1+\frac{X}{\tau_{D1}S^2})}}\right] + 1$$

where $N$ is the total number of particles, $f$ is the fractional contribution of the component, $X$ is time, $S$ is the structural parameter fixed to 5, $\tau_{D1}$ and $\tau_{D2}$ are the diffusion times of the two components, which are fixed to 27 $\mu$s and 126 $\mu$s, respectively, representing the diffusion time of free dye and Brichos. The resulting free Brichos particle number is shown in Figure 2**c**, and the average fraction of Brichos at the baseline or above the baseline (4% and 10%) is used to calculate the free Brichos concentration, which is fitted by the protein lipid binding model for stoichiometry.

We consider the 1:1 binding reaction: $\mathrm{P} + \mathrm{L} \underset{\mathrm{k_{off}}}{\overset{\mathrm{k_{on}}}{\rightleftharpoons}} \mathrm{PL}$, where our ligand $L$ is Brichos and our protein $P$ is a secondary nucleation site. At binding equilibrium, the total ligand concentration $c_L$ is:

$$c_L = [L] + [PL] = [L] + \frac{c_P[L]}{[L] + K_D}, \tag{15}$$

where $[L]$ is the concentration of free Brichos, $[PL]$ is the concentration of Brichos bound to secondary nucleation sites, $c_P$ is the total concentration of secondary nucleation sites, and $K_D$ is the dissociation constant. This can be rearranged to:

$$[L]^2 + [L](K_d + c_P - c_L) - c_L K_D = 0. \tag{16}$$

Finally:

$$[L] = -0.5 \cdot (K_d + M_0 s - c_L) + \sqrt{0.25 \cdot (K_D + M_0 s - c_L)^2 + c_L K_D}, \tag{17}$$

where $s$ is the number of monomer residues per secondary nucleation sites, and $M_0$ is the fibril mass concentration.

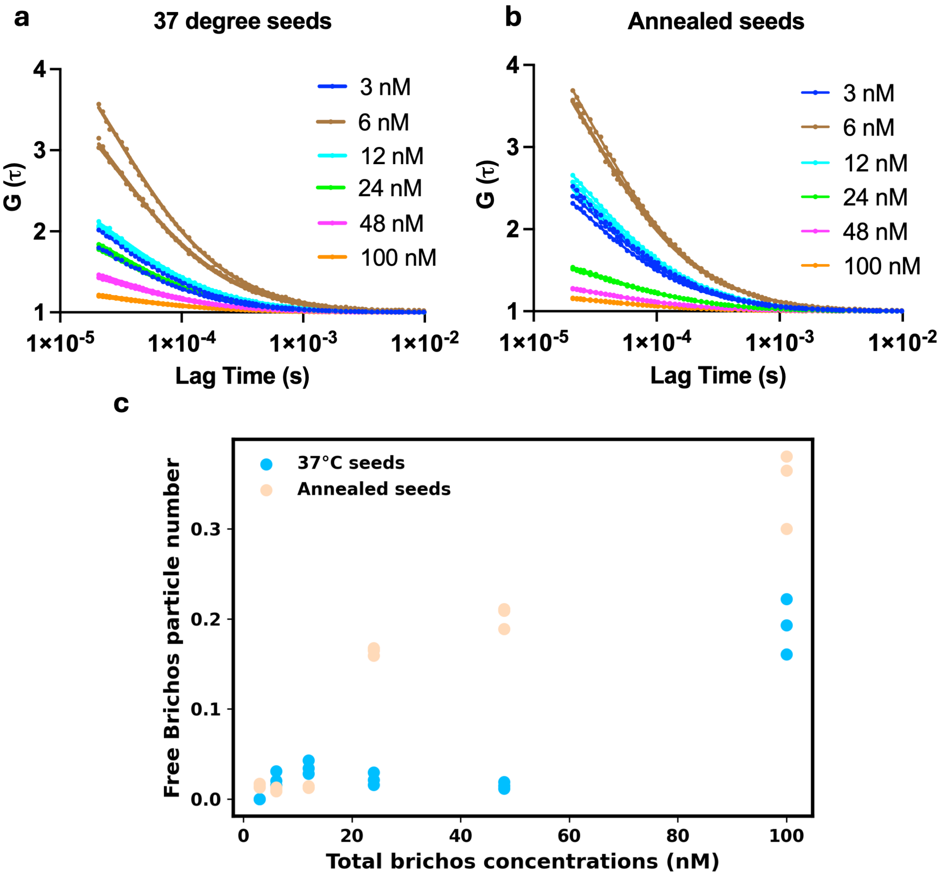


**Fig. 2** FCS experments of Aβ40 fibrils and Brichos. **a** and **b**: FCS fitting of the data for samples containing 4.5 $\mu$M 37°C fibrils or annealed fibrils, respectively. The fitting is done with 2 parameters fitting model while fixing the diffusion time of 27 $\mu$s (free dye) and 126 $\mu$s (Alexa488-Brichos). **c**: Free Brichos particle number gained from fitting in (a) and (b) is plotted against the total Brichos concentration.

### 4 NMR for solubility

The aggregation of 101 $\mu$M Aβ40 at 60℃ is monitored by integration of Aβ40 $^1$H NMR spectra on the methyl group region that is captured once per hour for 24 hours. NMR experiments were performed on a Bruker Avance III HD 900 spectrometer (Bruker Biospin, Rheinstetten, Germany), operating at a $^1$H resonance frequency of 899.8 MHz and fitted with a 5 mm cold probe.

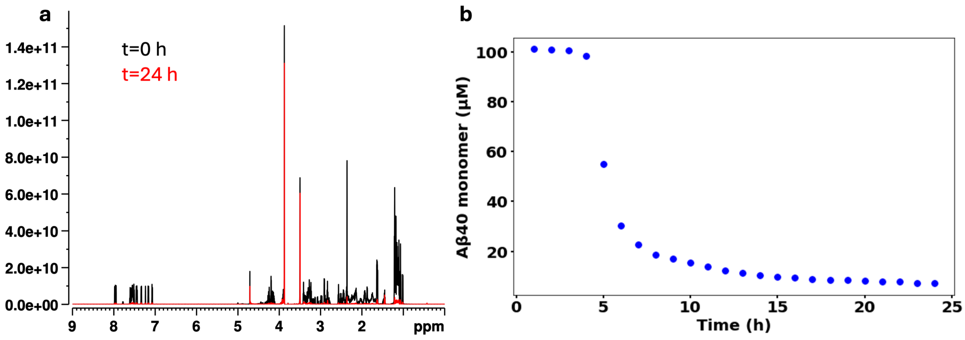


**Fig. 3** NMR experments of Aβ40 at 60℃. **a**: The $^1$H NMR spectra of 101 $\mu$M Aβ40 at 37℃ at t=0 hour (black) or t=24 hours (red). **b**: Aβ40 monomer concentration plotted against the time of incubation. The Aβ40 monomer concentration is calculated through integration of the methyl group peaks (1.0-1.4 ppm).

### 5 Static light scattering

The aggregation reaction under changing temperature was monitored in quartz cuvettes using a Probe Drum instrument (Probation Labs) by recording static light scattering at 5-minute intervals over up to 24 hours. The static light scattering signal was obtained by measuring the intensity of the excitation red laser at a 90° angle to the incident path. The 37℃ seeds were generated by incubating 10 µM A$\beta$40 with 2% seeds at 37℃ in the Probe Drum for at least 24 hours. The annealed seeds were produced by incubating 10 µM A$\beta$40 with 2% seeds, gradually cooling from 60℃ to 37℃ at a rate of 1℃ per hour, and then storing them at 37℃ until further experiments.

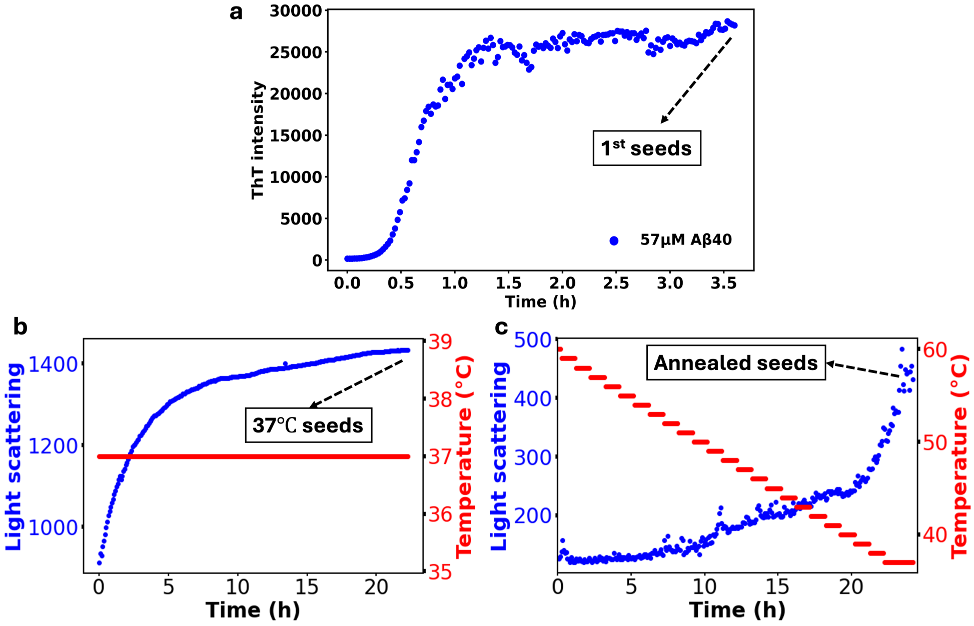


**Fig. 4** Generation of 37℃ fibrils or annealed fibrils. **a**: The aggregation of 57 $\mu$M Aβ40 at 37℃ is monitored by tracing the ThT (10 $\mu$M) intensity. A sample in the other well of the plate without ThT was incubated at the same time, and was taken at the end of aggregation as first seeds for generating 37℃ fibrils or annealed fibrils. **b** and **c**: The aggregation of 10 $\mu$M Aβ40 with 2% first seeds at 37℃ or 60 to 37℃ is monitored by light scattering. The fibrils at the end of aggregation is collected as 37℃ seeds or annealed seeds. Note that the light scattering values do not precisely quantify fibril mass, since sedimented fibrils are not detectable.

### 6 Nonequilibrium growth defect stoichiometry

As derived above, when defects are the equilibrium dislocation defect stoichiometry is given by the exponential of the misalignment free energy penalty. This can be rewritten as:

$$p_{mis} = \frac{e^{-G_{\mathrm{def}}/RT}}{e^{-G_0/RT}} = \frac{K_{\mathrm{def}}}{K_0}, \tag{18}$$

where $G_{\mathrm{def}}$ is the free energy of defect formation and $G_0$ the free energy of normal or correct elongation, and $K_{\mathrm{def}}$ and $K_0$ the associated equilibrium constants.

For simplicity we consider a 1-dimensional fibril with one monomer per plane. Then the expression in terms of equilibrium constants can be expanded as:

$$p_{eq} = \frac{2k_{+,d}}{k_{\mathrm{off},d}} \frac{k_{\mathrm{off}}}{2k_+}. \tag{19}$$

Here, $k_+$ and $k_{\mathrm{off}}$ have the usual meanings of the elongation and depolymerization rate constants (for correctly bound monomers), and $k_{+,d}$ and $k_{\mathrm{off},d}$ are the elongation and depolymerization rate constants for misaligned monomers.

In the limit of high supersaturation, essentially no depolymerization occurs and the stoichiometry of dislocations becomes just the ratio of elongation rates. Calling the stoichiometry in this "kinetic" limit $p_{neq}$, this is then $p_{neq} = k_{+,d}/k_+$. Our intuition is that increased kinetic trapping causes increased defect stoichiometry, i.e. $p_{neq} > p_{eq}$. This is true when:

$$\frac{k_{+,d}}{k_+} > \frac{2k_{+,d}}{k_{\mathrm{off},d}} \frac{k_{\mathrm{off}}}{2k_+}. \tag{20}$$

In turn, this is true if and only if:

$$\frac{k_{\mathrm{off},d}}{k_{\mathrm{off}}} > 1, \tag{21}$$

i.e. misaligned monomers detach more rapidly from fibril ends than correctly aligned monomers. This would seem to be self-evidently true, since they are not only less strongly bound but also closer in structure to soluble monomeric protein.

### 7 Growth defects with corresponding Cryo-TEM images

Cryo-EM imaging of Aβ40 or Aβ42 fibrils was performed as described previously [4]. Samples were vitrified on glow-discharged lacey carbon grids (300 mesh) using a Leica EM GP plunge freezer at 20°C and 90% relative humidity, and frozen in liquid ethane at −184°C. Grids were transferred to a JEM 2200FS electron microscope using a Fischione 2550 cryo-holder, and imaged at 200 kV under low-dose conditions with a 10 eV energy filter. Zero-loss images were acquired using a TVIPS F416 camera and processed using SerialEM and ImageJ. Changes in the twist periodicity or crossover distance were occasionally observed within individual Aβ40 or Aβ42 fibrils, indicating that fibril morphology can vary during elongation. Such variations likely reflect structural imperfections arising during growth. Different morphologies in the same fibril must be separated by growth defects. Branching Aβ42 fibrils were also observed; branch points are also a type of growth defect as outlined in Fig. 1 of the main text.

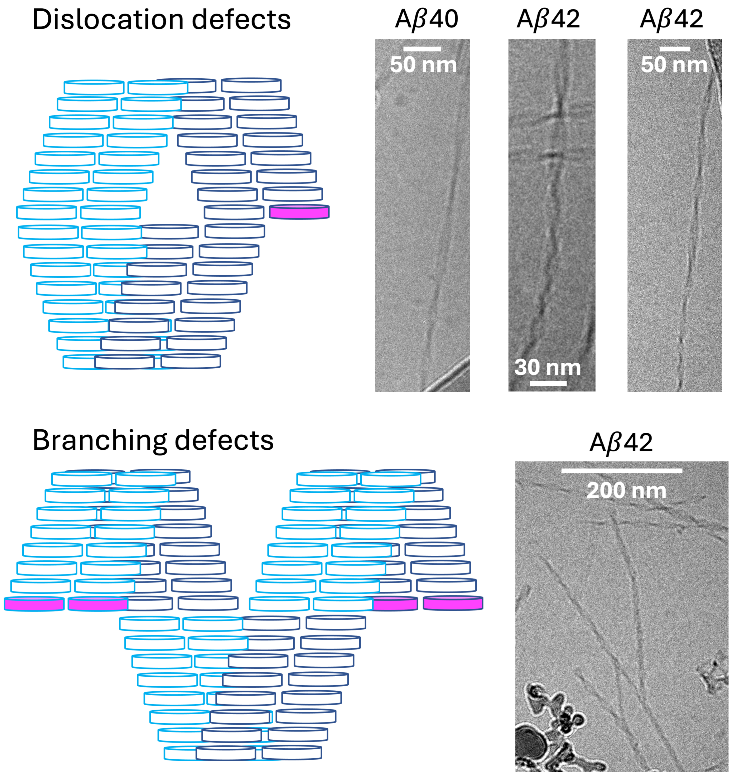


**Fig. 5** Cryo-TEM images of Aβ40 or Aβ42 fibrils showing evidence of growth defects. The Aβ40 fibrils are formed in a solution of 10 µM Aβ40 at pH 7.4, 37°C, while Aβ42 fibrils are formed in a solution of 5 µM Aβ42 at pH 8, 37°C.
